# Individual differences among deep neural network models

**DOI:** 10.1101/2020.01.08.898288

**Authors:** Johannes Mehrer, Courtney J. Spoerer, Nikolaus Kriegeskorte, Tim C. Kietzmann

## Abstract

Deep neural networks (DNNs) excel at visual recognition tasks and are increasingly used as a modelling framework for neural computations in the primate brain. However, each DNN instance, just like each individual brain, has a unique connectivity and representational profile. Here, we investigate individual differences among DNN instances that arise from varying only the random initialization of the network weights. Using representational similarity analysis, we demonstrate that this minimal change in initial conditions prior to training leads to substantial differences in intermediate and higher-level network representations, despite achieving indistinguishable network-level classification performance. We locate the origins of the effects in an under-constrained alignment of category exemplars, rather than a misalignment of category centroids. Furthermore, while network regularization can increase the consistency of learned representations, considerable differences remain. These results suggest that computational neuroscientists working with DNNs should base their inferences on multiple networks instances instead of single off-the-shelf networks.

## Introduction

Deep neural networks have recently moved into the focus of the computational neuroscience community. Having revolutionized computer vision with unprecedented task performance, the corresponding networks were soon tested for their ability to explain information processing in the brain. To date, task-optimized deep neural networks constitute the best model class for predicting activity across multiple regions of the primate visual cortex (Cadieu et al., 2014; Guclu and van Gerven, 2015; Khaligh-Razavi and Kriegeskorte, 2014; Schrimpf et al., 2018; Yamins et al., 2014). Yet, the advent of computer vision models in computational neuroscience raises the question in how far network internal representations generalize, or whether network instances, just like human brains, exhibit individual differences due to their distinct connectivity profiles. Strong differences would imply that the common practice of analyzing a single network instance is misguided and that groups of networks need to be analyzed to ensure the validity of insights gained.

Here we investigate individual differences among deep neural networks that arise from a minimal experimental intervention: changing the random seed of the network weights prior to training while keeping all other aspects identical. Our analyses of the network internal representations learned during training build on representational similarity analysis (RSA, Kriegeskorte, 2008), a multivariate analysis technique from systems neuroscience. RSA is based on the concept of representational dissimilarity matrices (RDMs), which characterize a system’s inner stimulus representations in terms of pairwise response differences. Together, the set of all possible pairwise comparisons provides an estimate of the geometric arrangement of the stimuli in high-dimensional activation space. The representations of two DNNs are considered similar if they emphasize the same distinctions among the stimuli, i.e. to the degree that their RDMs agree. Comparisons on the level of RDMs, which can be computed in source spaces of different dimensionality, thereby side-step the problem of defining a correspondence mapping between the units of the networks. Due to this, RSA is commonly used in cognitive computational neuroscience to compare DNNs to brain data. Comparing different DNN instances using the same technique therefore has the advantage that the current set of results will be directly applicable to the common neuroscientific use case. To quantify RDM agreement across network instances, we define *representational consistency* as the shared variance between network RDMs (squared Pearson correlation of the upper triangle of the RDMs; Figure 1).

**Fig 1.**
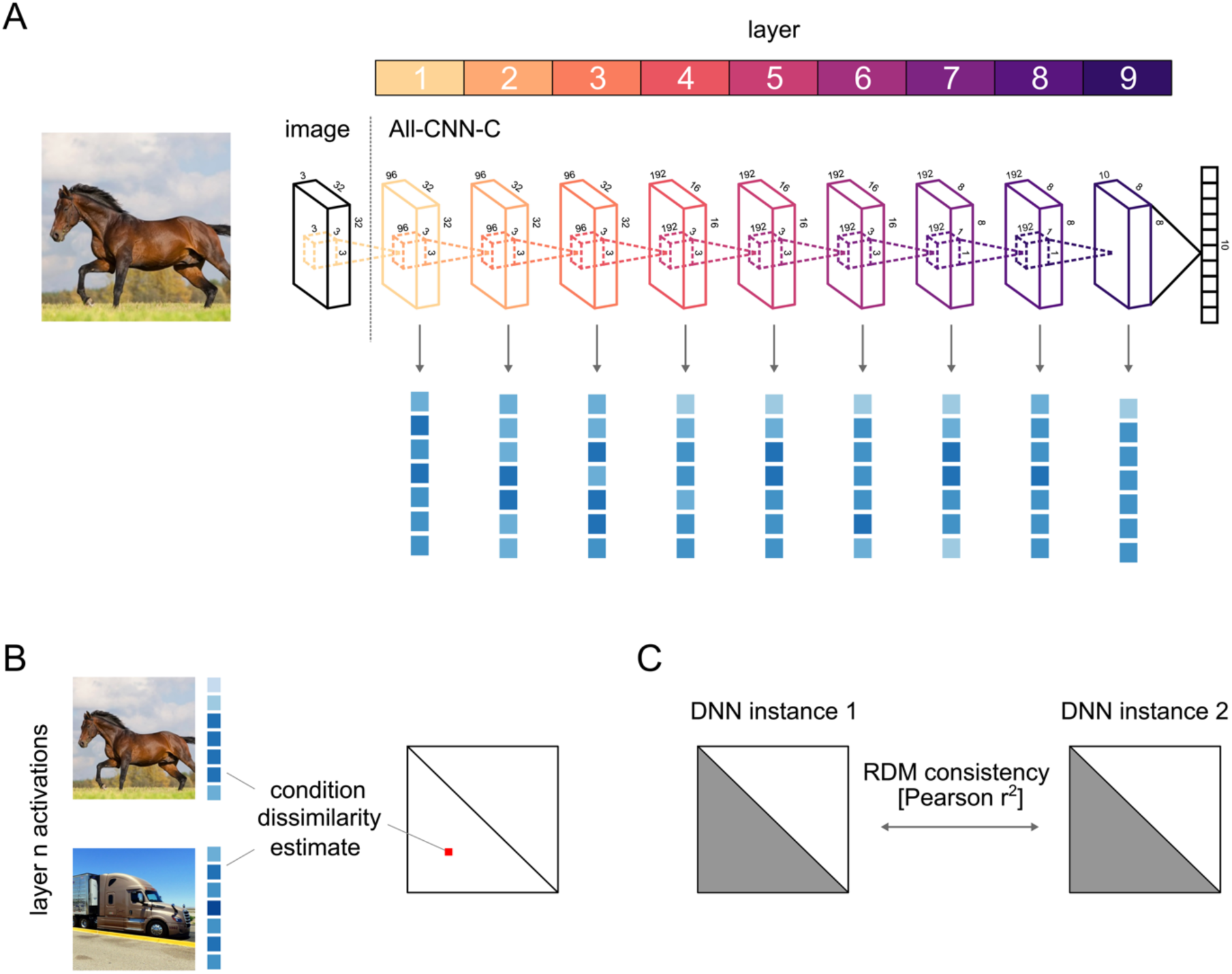
Characterizing network internal representations via representational similarity analysis and representational consistency. (**A**) Our comparisons of network internal representations were based on their multivariate activation patterns, extracted from each layer of each network instance as it responded to each of 1000 test images. (**B**) These high-dimensional activation vectors were then used to perform a representational similarity analysis (RSA). The fundamental building blocks of RSA are representational dissimilarity matrices (RDMs), which store all pairwise distances between the network’s responses to the set of test stimuli. Each test image elicits a multivariate population response in each of the network’s layers, which corresponds to a point in the respective high-dimensional activation space. The geometry of these points, captured in the RDM, provides insight into the nature of the representation, as it indicates which stimuli are grouped together, and which are separated. (**C**) To compare pairs of network instances, we compute their representational consistency, defined as the shared variance between network RDMs.

Based on this analysis approach, we visualize the internal network representations and test them for consistency. We then compare the size of the effects observed to differences between networks trained with different input statistics and test the reliability of the observations across multiple activity distance measures. Subsequently, we explore possible causes for these individual differences and investigate their interaction with network regularization.

## Results

We here investigate the extent to which deep neural networks exhibit individual differences. We approach this question by training multiple instances of the All-CNN-C network architecture (Springenberg et al., 2015) and a custom architecture (VGG-753) on an object classification task (CIFAR-10), followed by an in-depth analysis of resulting network internal representations. Network instances varied only in the initial random assignment of weights, while all other aspects of network training were kept identical. All networks performed similarly in terms of classification accuracy (ranging between 84.4 - 85.9% and 77.6 - 78.95% top-1 accuracy for All-CNN-C, and VGG-753, respectively).

To study and compare network internal representations, we extracted network activation patterns for 1000 test images (100 for each of the CIFAR-10 categories, Figure 1A) and characterized the underlying representations in terms of pairwise distances in the high-dimensional activation space (Figure 1B). The reasoning of this approach is that if two images are processed similarly in a given layer, then the distance between their activation vectors will be low, whereas images that elicit distinct patterns will have a large activation distance. The matrix of all pairwise distances (size 1000×1000) thereby describes the representational geometry of the test images, i.e. how exemplars of various object categories are grouped and separated by the units of a given layer (Kriegeskorte and Kievit, 2013).

### Stronger category clustering and individual differences in later network layers

To visualize the representational geometries of different network instances and layers, we projected the data into 2D using multidimensional scaling (MDS, metric stress). As can be seen in Figure 2 for two exemplary cases of All-CNN-C, subsequent network layers increasingly separate out the different image categories, in line with the training objective.

**Fig 2.**
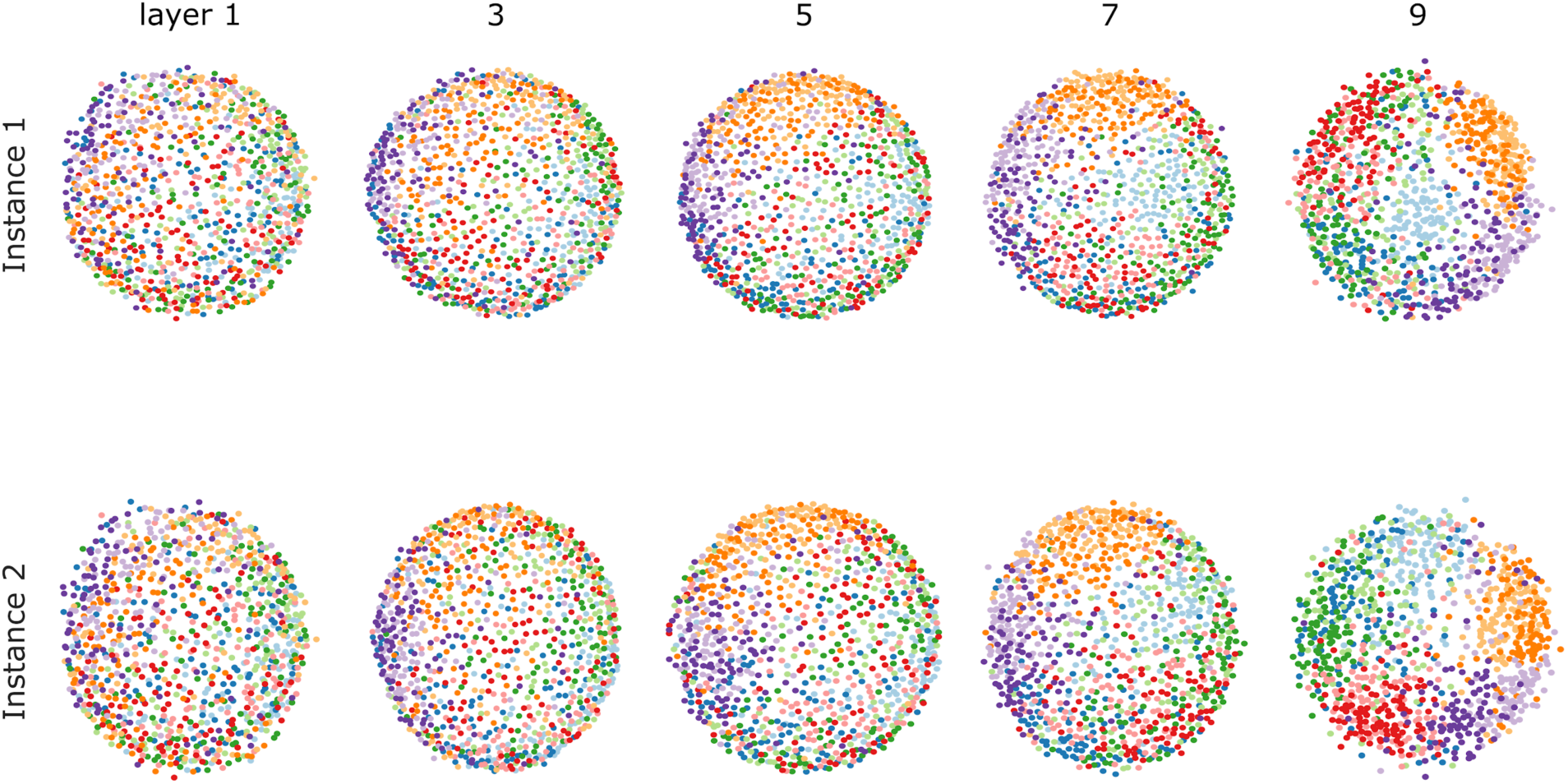
2D visualization of representational geometries in different depths of two network instances. The internal representations of two network instances were characterized based on their representational geometries. We computed the pairwise distances (correlation distance) between activity patterns in response to 1000 test stimuli from 10 visual categories and visualized them in 2D via multidimensional scaling (MDS; metric stress criterion). With increasing depth, networks exhibit increased category clustering and emerging differences.

Moving closer to the question of individual differences in network representations, we next investigated similarities on the level of RDMs. We again computed pairwise distances, but this time not based on activation patterns, but rather based on the network RDMs. Comparing patterns of representational distances has multiple benefits. For one, they offer a characterization of network internal representations that is largely invariant to rotations of the underlying high-dimensional space, including a random shuffle of network units (see Supporting Information for more details). Secondly, representational spaces of varying dimensionality can be directly compared, as the dimensionality of the RDM is fixed by the number of test images used.

This second-level distance measure was computed across all network layers and instances. Visualizing the respective distances in 2D (MDS, metric stress), we observe that representations diverge substantially with increasing network depth (Figure 3). While different network instances are highly similar in layer 1, indicating agreement in the underlying representations, subsequent layers diverge gradually with increasing network depth. Note that for later layers, the blue stripes parallel to the main diagonal indicate higher similarity across layers within a given network instance compared to the similarities across instances for a given network layer (see figure S2).

**Fig 3.**
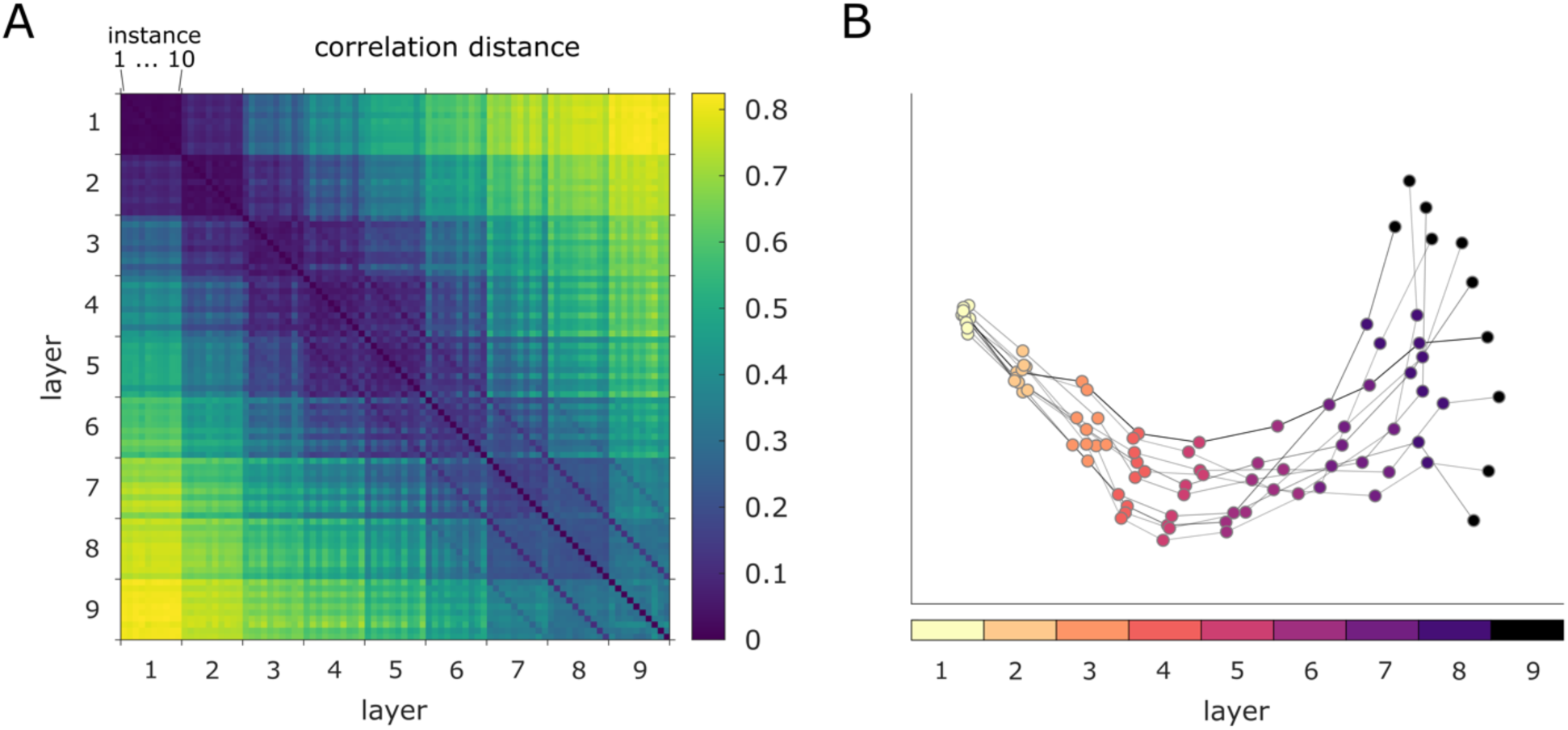
Network individual differences emerge with increasing network depth. (**A**) We compare the representational geometries across all network instances (10) and layers (9 convolutional) for All-CNN-C by computing all pairwise distances between the corresponding RDMs. (**B**) We projected the data points in (A) (one for each layer and instance) into 2D via MDS. Layers of individual network instances are connected via grey lines. While early representational geometries are highly similar, individual differences emerge gradually with increasing network depth.

### Representational consistency decreases with increasing network depth

Following this initial qualitative assessment, we performed quantitative analyses for each network layer by testing how well the distribution of representational distances generalizes across network instances. This was accomplished by computing *representational consistency*, defined as the shared variance between the lower triangle of the respective RDMs (Figure 1 C, each triangle contains 499,500 distance estimates, results are obtained from 45 pairwise network comparisons for each respective layer and network architecture as 10 network instances are trained for each architecture). This measure of consistency is based on all pairwise distances between category exemplars (100 exemplars for 10 categories each). We therefore refer to this as exemplar-based consistency.

Two network architectures were tested (All-CNN-C, and VGG-753, see methods for details). Correlation distance was chosen as dissimilarity measure in computing RDMs, as it is currently the most frequently used distance measure in systems and computational neuroscience. As shown in Figure 4, representational consistency drops substantially with increasing network depth for both network architectures. To get better insights into the size of this effect, additional networks were trained (i) based on different images originating from the same categories, and (ii) based on different categories (see methods for details). The observed drops in consistency for different weight initializations (to about 44% and 71% for All-CNN-C and VGG-753, respectively), are comparable to training the networks with the same distribution of categories but completely separate image datasets (Figure 5, blue vs. orange).

**Fig 4.**
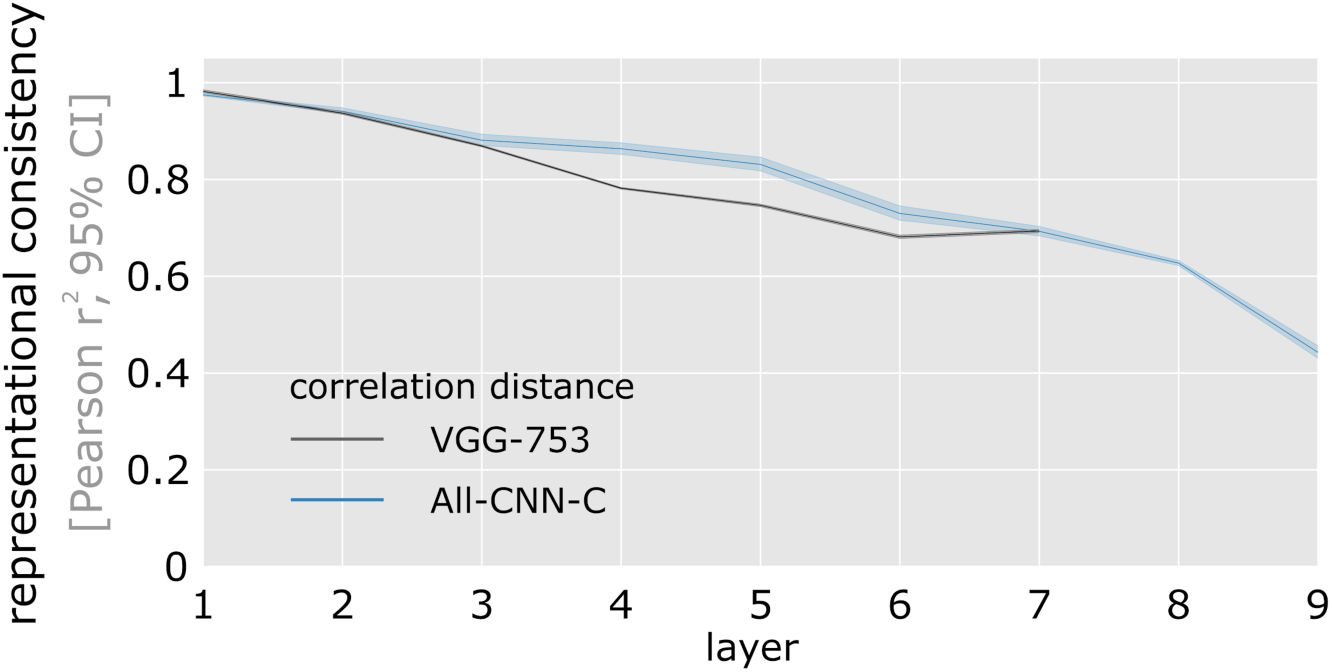
Representational consistency decreases with increasing network depth. Shown is the average representational consistency for each layer computed across all pairwise comparisons of network instances (45 comparisons for 10 instances, computed separately for two network architectures). Error bars indicate 95% confidence intervals (bootstrapped).

**Fig 5.**
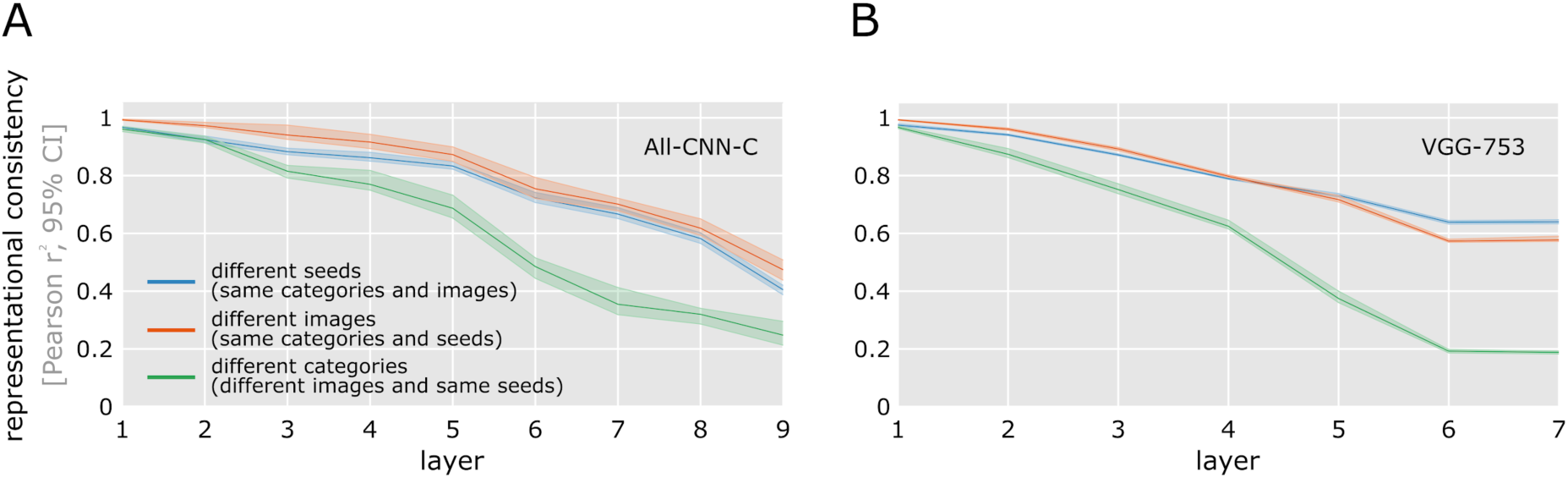
Representational consistency declines with increasing network depth when trained on separate image sets. To better understand the size of the effect in Figure 4, we trained a separate set of networks based on (**A**) All-CNN-C, (**B**) VGG-753) while using different images from the same categories but the same seeds (orange), and different categories, different images, and same seeds (green). The minimal intervention of using a different seed for the random weight initialization (shown in blue, data equivalent to Figure 4) affects the internal representations about as much as using a completely different set of training images (10 categories per training set; orange). Please note that part of the larger drop in representational consistency for training with different categories (5 categories per training set; green) can be attributed to training only five categories while computing the RDMs with images from all 10 categories.

To ensure that the effects observed are not specific to correlation distance used in computing the RDMs, additional analyses were performed based on cosine, (unit length pattern-based) Euclidean distance and norm difference (measuring the absolute difference in the norm activation vectors, Figure 6). In all cases, representational consistency was observed to drop considerably with increasing network depth. These results demonstrate that while different network instances reach very similar classification performance, they do so via distinct internal representations in the intermediate and higher network layers.

**Fig 6.**
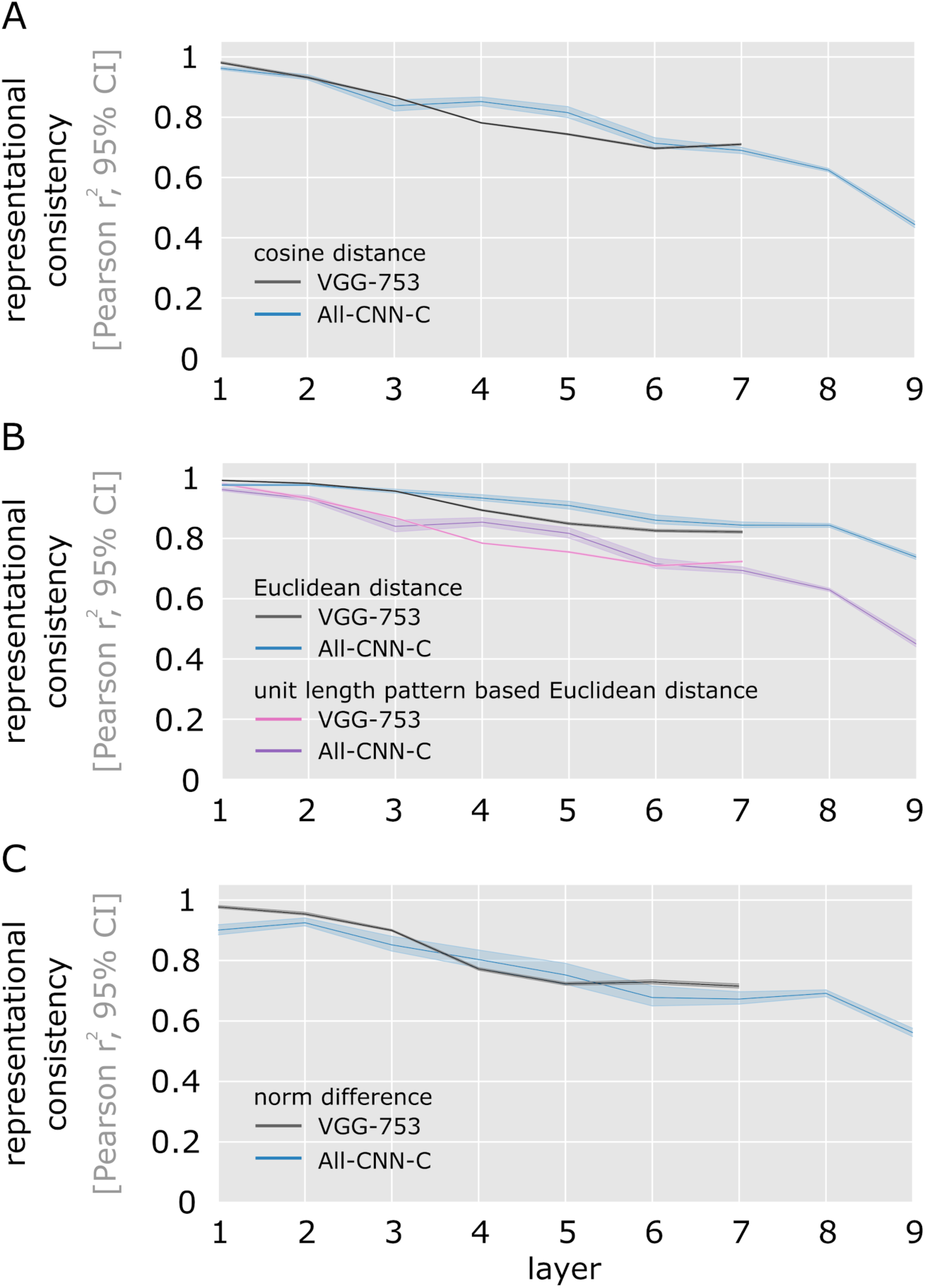
Representational consistency declines with increasing network depth irrespective of distance measure. Representational consistency decreases with increasing layer depth for both tested DNN architectures, and across multiple different ways to measure distances in multivariate population responses (cosine (**A**), Euclidean distance and unit length pattern-based Euclidean distance (**B**), and differences in vector norm (**C**)). We show the average representational consistency for each layer, computed across all pairwise comparisons of network instances (45 comparisons for 10 instances), together with a 95% bootstrapped confidence interval.

### Causes of decreasing representational consistency

We have shown above that different network instances can exhibit substantial individual differences in their internal representations. Next, we investigated potential mechanisms that may contribute to this effect.

Our first analyses are based on the observation that the training goal of maximal category separability does not put a strong constraint on the relative positions of categories and category exemplars in high-dimensional activation space. To investigate this possibility, for the 10 network instances of All-CNN-C used in the previous section, we first computed a category clustering index (CCI) for each network layer using the network responses to the set of 1000 test images (drawn from 10 categories). CCI is defined as the normalized difference in average distances for stimulus pairs from different categories (across) and stimulus pairs from the same category (within): CCI = (across - within) / (across + within). CCI approaches zero with no categorical organization and is positive if stimuli from the same category cluster together (maximum possible CCI = 1). We find a negative relationship between CCI and representational consistency (Pearson r = −0.92, p = 0.001; (Pernet et al., 2013)), indicating that network layers that separate categories better do exhibit stronger individual differences.

This correlation is consistent with two possible scenarios: networks can exhibit a different arrangement of the overall category clusters, and different arrangements of individual images within the category clusters, as both are not constrained by the training objective to categorize. To investigate the variability in general cluster placement, we computed representational consistency based on the ten category centroids (RDMs computed from the pairwise distances of average response patterns for each category). This analysis revealed that this centroid consistency is considerably higher than the previous exemplar-based consistency (Figure 8A, μ_centroid-based_ = 0.8801, CI_95_ = [0.8700, 0.8905] vs. μ_exemplar-based_ = 0.4429, CI_95_ = [0.4291, 0.4551] for correlation distance; μ_centroid-based_ = 0.9515, CI_95_ = [0.9450, 0.9571] vs. μe_exemplar_based_ = 0.7384, CI_95_ = [0.7312, 0.7466] for Euclidean distance, all computed for the final layer of All-CNN-C). This finding cannot be explained by the lower number of pairwise comparisons (45 vs. 499,500 for centroid and stimulus RDMs, respectively) or the operation of averaging large numbers of activation patterns (each centroid is computed based on 100 activation patterns), as computing centroids from random stimulus assignments yielded significantly lower centroid-based representational consistency (95% CI of centroid-based consistency based on random class assignment [0.14, 0.81], Figure 8B). The reliable arrangement of category centroids suggests that the main source of the observed individual differences lies in the arrangement of category exemplars within the category clusters. This view was corroborated by computing consistency not on the whole exemplar-based RDM that contains all pairwise distances, but only on the dissimilarities of exemplars of the same categories (within-category consistency). Here we observe a drop in consistency that is largely comparable to the original decrease for exemplar-based consistency when investigating the whole RDM (Figure 8A).

**Fig 8.**
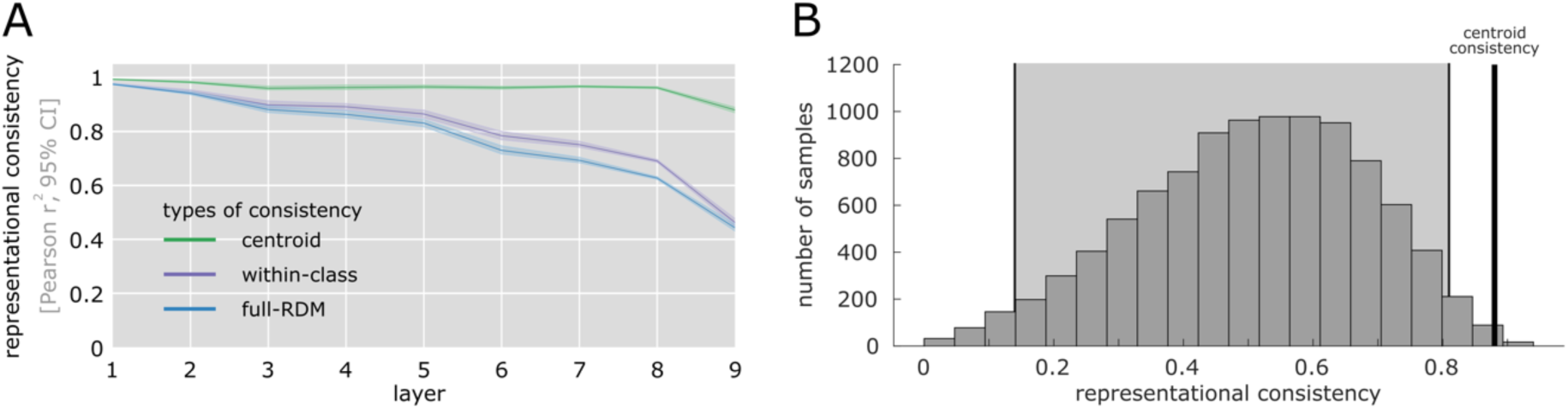
Category centroids are highly consistent across network instances. (**A**) Centroid-based representational consistency (green) remains comparably high throughout, whereas the consistency of within-category distances decreases significantly with increasing network depth. This indicates that differences in the arrangement of individual category exemplars, rather than large scale differences between class centroids are the main contributor to the observed individual differences. (**B**) High centroid-based representational consistency cannot be explained by the smaller RDMs or the averaging of multiple response patterns, as centroids of randomly sampled classes show a significantly lower mean consistency (95% CI in light grey background).

In addition to an individual placement of category centroids and category exemplars, some properties of the underlying dissimilarity measures can be a source for lower representational consistency, especially in cases of a rotated representational space. Many commonly used DNNs use rectified linear units (ReLUs) as a nonlinear operation, resulting in unit activations ≥ 0. While overall rotations of this all-positive space will not affect classification performance, they can affect correlation and cosine distances (see Figure 9, and Figure S1 demonstrating the additional effect that rotations around the origin affect correlation distances but not cosine distances).

**Fig 9.**
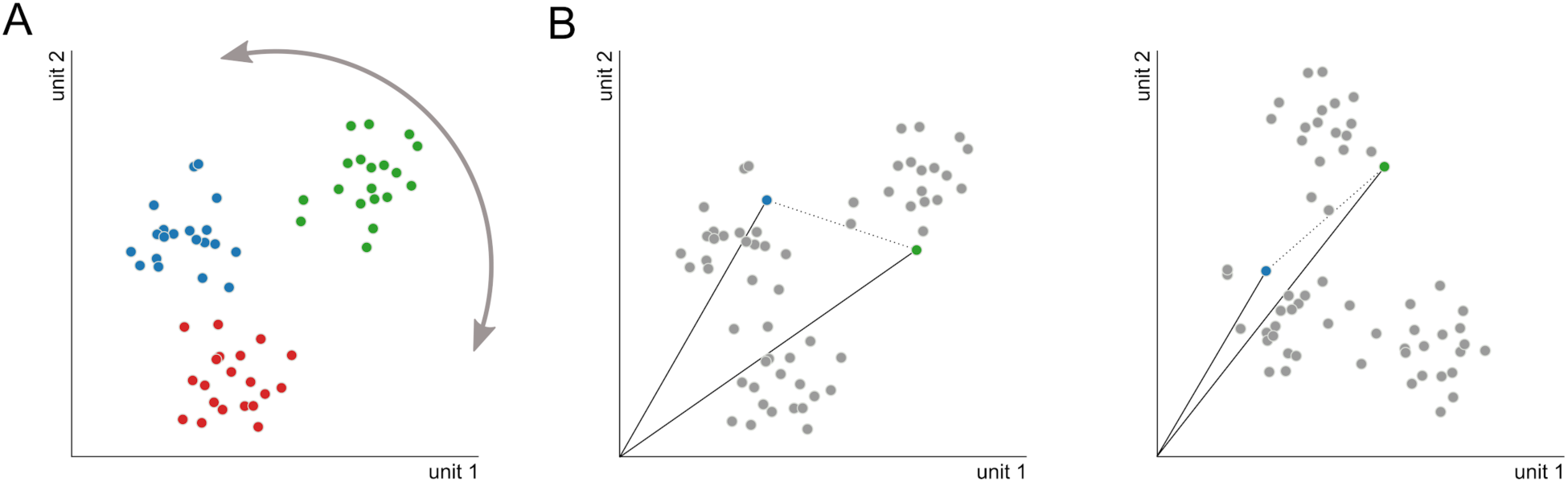
Rotation of a ReLU activation space affects correlation- and cosine-distance estimates. (**A**) Three exemplary classes (blue, green, red) are rotated in the all-positive (post-ReLU) activation space, here shown as a 2D example. (**B**) When comparing the activation space before (**left panel**) and after the rotation, the angle between pairs of images can differ markedly, thereby leading to lower representational consistency despite an overall stable data arrangement. (see Figure S1 for simulations using correlation distance).

To test the magnitude of this effect, we subtracted the mean activation pattern across all test images from the units of a given layer (cocktail blank normalization). As shown in Figure 10, this normalization leads to increases in representational consistency for RDMs computed using correlation or cosine distance. While the size of the effect is comparably small, these results indicate that a cocktail blank normalization can be of potential benefit when comparing correlation- or cosine-based RDMs of multiple DNNs or DNNs and brain data.

**Fig 10.**
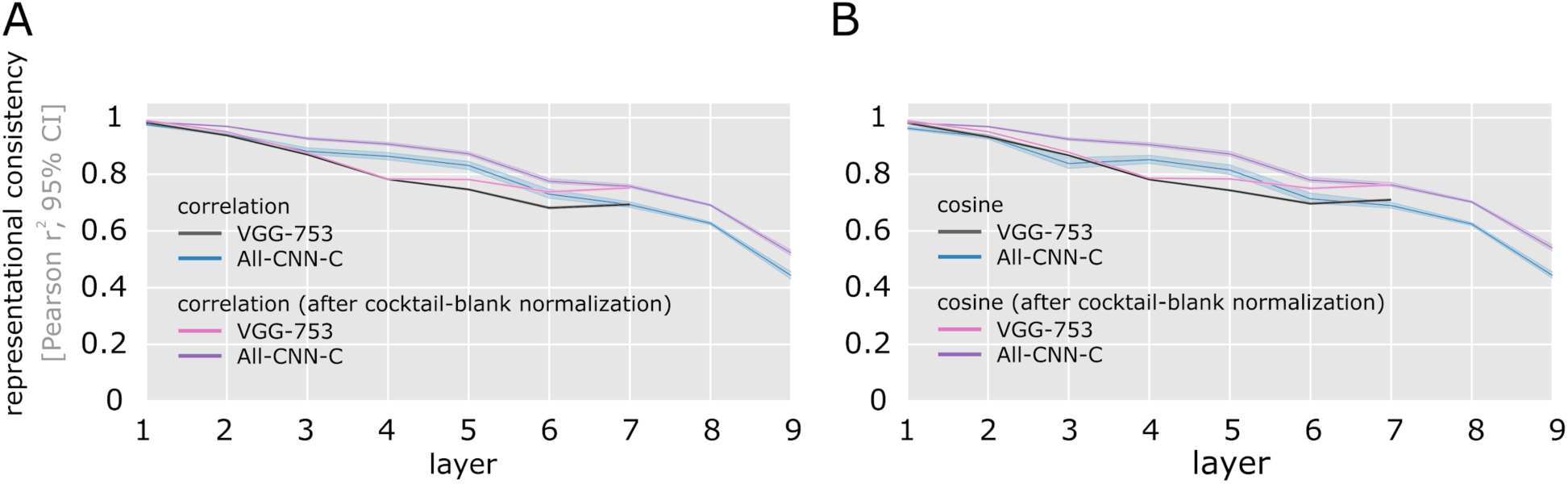
Cocktail blank normalization increases consistency for correlation and cosine distances. Centring the data via cocktail blank normalization increases representational consistency for correlation (**A**) and cosine distance (**B**). Euclidean distance measures are not affected, as the resulting representational geometries are rotationally invariant.

### Network regularization (Bernoulli dropout) affects representational consistency

An explanation of individual differences via missing constraints imposed by the training objective raises the possibility that explicit regularization during network training can provide the missing representational constraints (McClure and Kriegeskorte, 2018; Srivastava et al., 2014). We investigated this possibility experimentally by training networks at various levels of dropout regularization. We trained 10 network instances of All-CNN-C for each of 9 dropout levels (Bernoulli dropout probability ranging from 0 to 0.8, a total of 90 network instances trained) and subsequently tested the resulting representations for their ability to classify input, and for their representational consistency. To test for differences in task performance, we computed the top-1 categorization accuracy for the training- and test data. For the test data, we contrast network inference with and without dropout. In line with the literature (Srivastava et al., 2014), we find reduced training accuracy, but enhanced test accuracy at moderate dropout levels (Figure 11 A).

**Fig 11.**
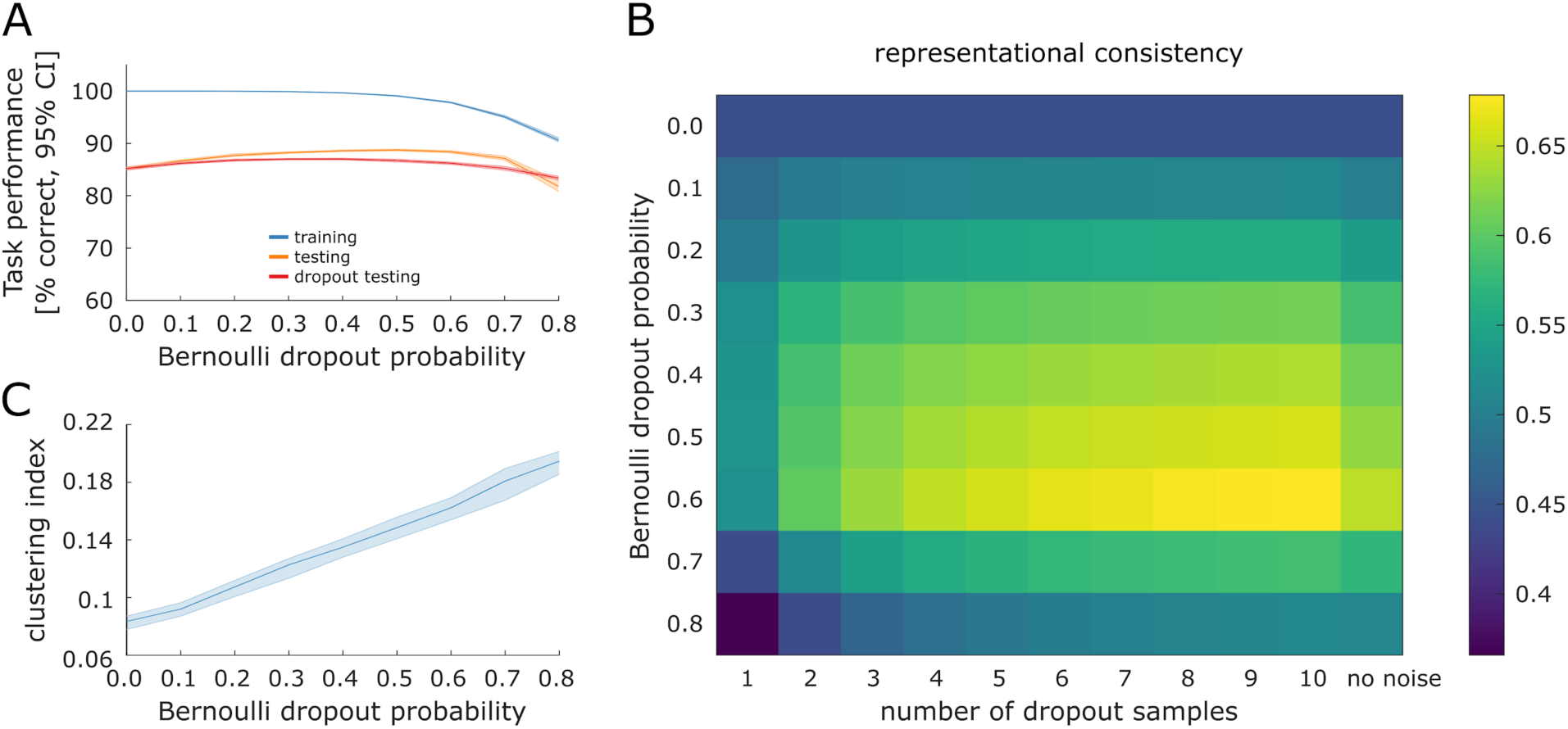
Effects of dropout regularization on task performance and representational consistency. (**A**) Average task performance across all network instances (shown with 95% CI) for the training set (blue), test set (orange), and when using dropout sampling at inference time for the test set (red, 1 sample). (**B**) Representational consistency in the final convolutional layer of All-CNN-C as a function of dropout probability during training and test (dropout probability at test time equal to training dropout). When using dropout at test time, multiple samples can be drawn for each stimulus in the test set (creating multiple RDMs). Consistency for network pairs was computed for the respective average RDM for each instance. Consistency was observed to be highest when 10 samples were obtained from a DNN trained and tested at a dropout rate of 60%. (**C**) The clustering index (also see Figure 7) increases with increasing Bernoulli dropout probability, here shown for the penultimate layer of All-CNN-C.

The effects of dropout training on representational consistency were again investigated using layer 9 of All-CNN-C, which exhibited the lowest consistency levels in our original analyses. These analyses revealed that dropout regularization yields increased representational consistency across network instances. When using no dropout at test time, a dropout probability of 0.6 during training provides the highest consistency level, reaching an average of 64.7% shared variance (rightmost column in Figure 11 B).

In analogy to our analyses of test accuracy when applying dropout at the time of inference, we investigated in how far this may affect representational consistency estimates. For each network instance, we computed 10 RDM samples while keeping the dropout mask identical across network instances and the rate identical to training. The average of a varying number of up to 10 RDM samples was subsequently used to compute representational consistency across network instances. We find that increasing the number of RDM samples led to increased representational consistency for all dropout levels. Maximum representational consistency was observed for 10 RDM samples at a dropout probability of 0.6, reaching an average of 67.8% shared variance across network instances. This suggests that dropout applied during training and test can increase the consistency of the representational distances across network instances.

As a possible explanation for how dropout could have affected representational consistency, we computed the category clustering index (CCI) for the penultimate layer of All-CNN-C and different dropout levels. This is based on the idea that stronger clustering around the category centroids in the latest network layer will at the same time yield higher consistency, as the arrangement of category centroids is highly consistent. As shown in Figure 11C), we observe a positive relationship between dropout probability and category clustering. However, while clustering is further enhanced for dropout levels >0.6, representational consistency starts decreasing. To further explore this effect, we re-computed centroid consistency for highest dropout level (0.8) and observed that centroid consistency is significantly decreased (μ_dropout=0.8_ = 0.7422, CI_95_ = [0.6881, 0.7854]) compared to the no dropout case (μ_no_dropout_ = 0.8801, CI_95_ = [0.8700, 0.8905]). Thus, while denser clustering around centroids increases consistency in cases where the centroids themselves are consistent, high levels of dropout lead to less consistent centroids and therefore to an overall decrease in consistency.

### Representational consistency across training trajectories

We have observed above that representational consistency across network instances is remarkably stable for category centroids. This raises the question as to whether this alignment is the result of task training, or whether category centroids are already well-aligned early during training. To investigate this, we computed representational consistency (exemplar-based and centroid-based) across different network instances and training epochs. We extracted activation patterns from each network instance at different stages of training and subsequently computed pairwise representational consistency, again using the penultimate layer of All-CNN-C. Individual networks exhibit high consistency after the first epoch, which however decreases from thereon, indicating that task training enhances individual differences. Yet, from very few epochs onwards networks exhibit stable representations with each network remaining on its own learning trajectory (Figure 12A, multiple diagonal lines indicate stable representations across training compared to other network instances). Consistency seems to saturate from epoch 150 onwards, indicating overall smaller changes in the network internal representations. Consistent with our earlier results, centroid-based consistency is overall higher across network instances even for the earliest epochs (Figure 12B). These results indicate that task training leads to decreased consistency, whereas learning trajectories of individual networks across time remain surprisingly robust.

**Fig 12.**
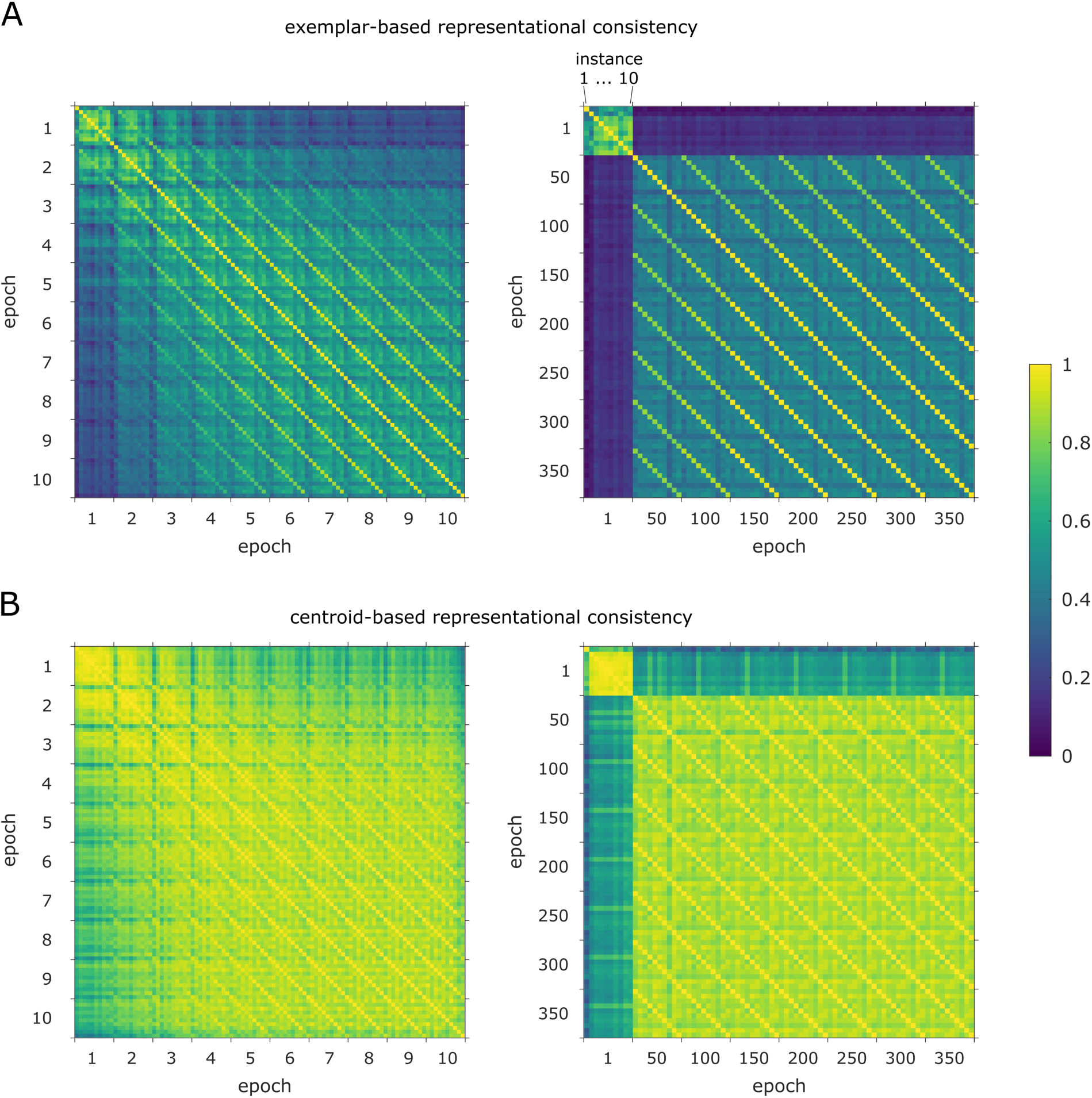
Penultimate-layer representational consistency across training consistency for RDMs based on individual images and on class centroids. (**A**) Exemplar-based representational consistency across epochs [1 to 10] (left) and across epochs [1 to 350 in steps of 50] (right). (**B**) Same as **A**, but RDMs are based on class centroids instead of individual images. After the first epoch the representations across network instances show elevated image- and centroid-based consistency (left panel in both **A** and **B**, respectively). However, consistency decreases with subsequent epochs, indicating that task training increases individual representational differences. From very few epochs onwards, representations become remarkably stable, saturating around epoch 150, indicating comparably smaller representational adjustments induced by training.

## Discussion

In a series of experiments, we here investigated how the minimal intervention of changing the initial set of weights in feedforward deep neural networks, while keeping all else constant, affects their internal representations. Importantly, while many metrics exist to compare DNN representations (Kornblith et al., 2019; Raghu et al., 2017; Wang et al., 2018), our current analyses were explicitly chosen to match techniques commonly used in neuroscience to compare DNNs to brain data. The current set of results therefore directly transfers to the relevant neuroscientific publications. Operationalized as representational consistency, we demonstrated that significant individual differences emerge with increasing layer depth. This finding held true for various distance measures commonly used to compute the RDMs (correlation distance, cosine distance, variants of Euclidean distance, and norm differences). RDMs computed from Euclidean distances showed the least differences. In part, this can be attributed to the fact that this distance measure is sensitive to differences in overall network activation magnitudes, which may overshadow more nuanced pattern dissimilarities, in line with the lower consistency observed for norm-standardizing Euclidean distances (unit-length pattern based Euclidean distance). Although further experiments are required, we expect our results to generalize to representations learned by (unrolled) recurrent neural network architectures (Kar et al., 2019; Spoerer et al., 2019), if not explicitly constrained (Kietzmann et al., 2019b). For an investigation of recurrent neural network dynamics arising from various network architectures see Maheswaranathan et al. (2019).

Having demonstrated significant network individual differences, we explored multiple non-exclusive explanations fort the effects. Based on the hypothesis that the network training objective of optimizing for categorization performance may not sufficiently constrain the arrangement of categories and individual category exemplars, we analyzed category clustering, centroid arrangement, and within-category dissimilarities. All of these analyses point to a high consistency of category centroids, rendering differences between individual category exemplars the main contributor of the differences observed. As an additional source of variation, we identified an interaction between properties of the distance measures used and the ReLU nonlinearity in the DNNs. We showed that cocktail blank normalization in the DNN activation patterns can increase consistency for measures that are not robust to rotations that are not centered around zero (cosine distance) or general rotations (correlation distance). In addition to this, we showed that network regularization via dropout during training and test can enhance representational consistency estimates. As a partial explanation for this increase, we demonstrated that category centroids are highly consistent and that dropout enhances category clustering.

Our finding of considerable individual differences has implications for computational neuroscience where single pre-trained computer vision networks are often used as models of information processing in the brain. Neglecting the potentially large variability in network representations will likely limit the generality of claims that can be derived from comparisons between DNNs and neural representations. While we here present multiple approaches that can increase consistency (cocktail-blank, dropout, and the choice of distance measure), significant differences remain. For computational neuroscience to take full advantage of the deep learning framework (Cichy and Kaiser, 2019; Kietzmann et al., 2019a; Kriegeskorte and Douglas, 2018; Richards et al., 2019), we therefore suggest that DNNs should be treated similarly to experimental participants, as analyses should be based on groups of network instances. Representational consistency as defined here will give researchers a way to estimate the expected network variability for a given training scenario, and thereby enable them to better estimate how many networks are required to ensure that the insights drawn from them will generalize. In addition to the impact on computational neuroscience, we expect the concept of representational consistency, which can be applied across different network layers, architectures, or training epochs, to also benefit machine learning researchers in understanding differences among networks operating at different levels of task performance.

## Materials and methods

### Deep neural network training

The main architecture used throughout all experiments presented here is All-CNN-C (Springenberg et al., 2015), a 9 layer fully convolutional network that exhibits state of the art performance on the CIFAR-10 dataset. To optimize architectural simplicity, the network uses only convolutional layers with a stride of 2 at layer 3 and 6 to replace max- or mean-pooling. We used the same number of feature maps (96, 96, 96, 192, 192, 192, 192, 192, 10) and kernel-sizes (3, 3, 3, 3, 3, 3, 3, 1, 1) as in the original paper (Figure 1 A).

To show that our results generalize beyond a single DNN architecture we trained an additional architecture reminiscent of VGG-S (Chatfield et al., 2014). In contrast to the original VGG-S architecture, we replaced the two deepest, fully-connected layers with convolutional layers to reduce the number of trainable parameters and thus the training duration by ∼80%. The number of feature maps used per layer was [96, 128, 256, 512, 512, 1024, 1024], and the kernel sizes were [7, 5, 3, 3, 3, 3, 3]. We used ReLU as the activation function at every layer. Mirroring the kernel sizes across layers, we refer to this architecture as “VGG-753”.

All-CNN-C network instances were trained for 350 epochs using a Momentum term of 0.9 and a batch size of 128. All networks of the VGG-753 architecture were trained for 250 epochs using ADAM with an epsilon term of 0.1 and a batch size of 512. For both architectures, we used an initial learning rate of 0.01, the L2 coefficient was set to 10^−5^, and we performed norm-clipping of the gradients at 500. Training of the main DNNs was performed on the full CIFAR-10 image set. CIFAR-10 consists of 10 categories of objects, each of which is represented by 5,000 training and 1,000 test images. Ten network instances were trained for the main analyses, all without dropout.

Network training was identical across all instances (same architecture, same dataset, same sequence of data points), with the exception of the random seed for the weight initialization. As a result, the networks only differ in the initial random weights, which are, however, sampled from the same distribution (He et al., 2015).

### Comparing layer-internal representations across network instances

#### Representational similarity analysis and representational consistency

We characterize the internal representations of the trained networks based on representational similarity analysis (RSA, Kriegeskorte, 2008), a method used widely across systems neuroscience to gain insight into representations in high-dimensional spaces.

RSA builds upon the concept of representational dissimilarity matrices (RDMs), which store all pairwise distances between the stimulus-driven pattern activations in response to a large set of input stimuli (Figure 1 A). Here we use 1,000 test stimuli, 100 from each of the 10 CIFAR-10 categories, such that the resulting RDMs have a size of 1000×1000 (Figure 1 B). The RDMs are symmetric around the diagonal and therefore contain 499,500 unique distance estimates. In the current set of experiments, pairwise distances (using correlation-, cosine-, and (unit length pattern-based) Euclidean-distance) are measured in the activation space of individual layers, where each unit corresponds to its own input dimension. The resulting matrix thereby characterizes the representational space spanned by the network units, as it depicts the geometric relations of all different input stimuli with respect to each other. This focus on relative distances renders RSA largely invariant to rotations of the input space (including random shuffling of input dimensions, but see Figure S1). It is therefore well suited for comparisons across deep neural network instances.

Because RDMs are distance matrices, they can be used as a basis for multidimensional scaling (MDS) to project the high-dimensional network activation patterns into 2D. While not a lossless operation, as high-dimensional distances can usually not be perfectly reproduced in 2D, MDS does nevertheless enable us to gain first insights into the internal organization by visualizing how network layers cluster the 1000 test images from the 10 different categories.

In addition to enabling 2D visualizations of network internal representations (or, put differently, the organization of test-images in high-dimensional layer activation space, Figure 2), RDMs themselves can be used as observations (each RDM is a point in the high-dimensional space of all possible RDMs) and thereby form the basis for computing “second-level” distance matrices. The resulting distance matrices can be used to compare representations across multiple network layers and network instances (rather than test-images as in first-level RDMs). Here, we compute a second level distance matrix based on the RDMs for all network layers and instances. Again, we use MDS to visualize the data points in 2D (Figure 3).

For a more quantitative comparison of network internal representations, characterized here in terms of RDMs, we define **representational consistency** as the shared variance across representational distances observed in high-dimensional network activation space. Representational consistency is computed as squared Pearson correlation between RDMs (Figure 1 C). If two network instances separate the test stimuli with similar geometry, the representational consistency will be high (max 1), whereas uncorrelated RDMs exhibit low representational consistency (min 0).

#### Comparing the effect of weight initialization to the effects of varying input statistics

The main experimental manipulation in this work consists of using different random weights at the point of network initialization. To better understand the size of the effects on network internal representations, we compared the effects observed to differences that emerge from using different images from the same categories (within-category split), or different categories altogether (across-category split). To perform this control analysis, two subsets of CIFAR-10 were created. For the across-category division, we split the training and test sets on the level of categories. This resulted in two datasets with 5 categories each while preserving the number of images per category (5,000 training, 1,000 test images). For the within-category division, the dataset was split based on images rather than categories. This preserves the number of categories (10) but halves the number of training images per category. For an illustration of the splitting procedure that resulted in the within-category, and the across-category splits of CIFAR-10, see Figure 13.

**Fig 13.**
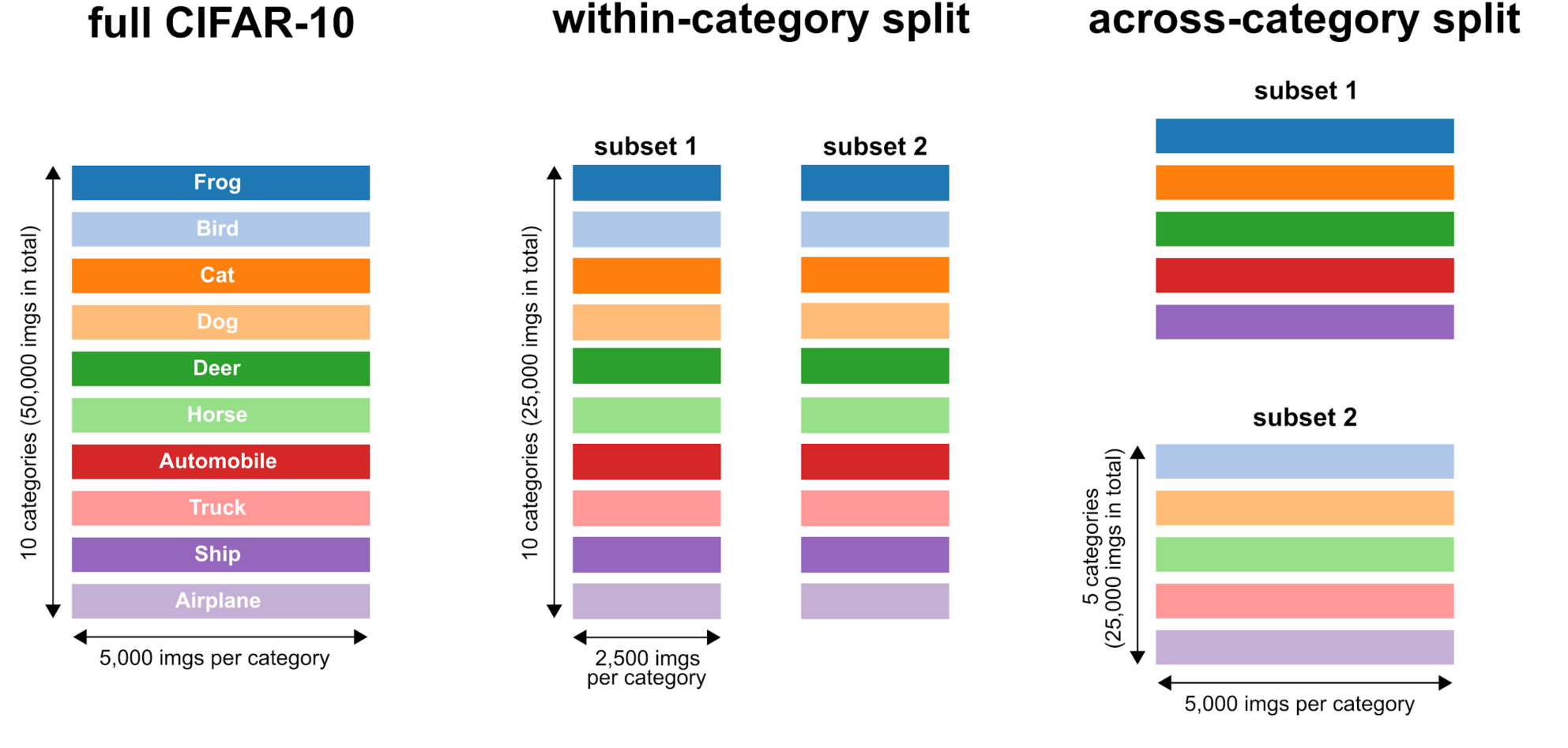
Visualization of the CIFAR-10 training sets used. Different categories are shown as distinct colors. Left panel: The full CIFAR-10 training set consists of 10 categories with 5,000 images each, 50,000 images in total. **Center panel**: the within-category split dataset contains 10 categories with 2,500 images each, 25,000 images in total for each subset. **Right panel**: the across-category split dataset contains 5 categories with 5,000 images each, again 25,000 images in total for each subset. When splitting across categories, the number of animal- and vehicle-categories of the full CIFAR-10 set was equally distributed across the two subsets.

In summary, the consistency of network instances resulting from different random weight initializations (different seeds, same categories, same images), was compared with (a) different images (same seed, same categories), and (b) different categories (same seed, different images; Figure 5). Five networks were trained for each half of the dataset for both splits (a, and b, resulting in 5×2=10 network instances each). Representational consistency was computed using pairs of network instances with the same random seed (5 pairs for each split). Note that representational consistency was computed based on 1,000 test images from all 10 CIFAR-10 categories, independent of the training set used to train the networks.

#### Category clustering and its relation to representational consistency

To measure how well the layers of a network separate exemplars from different categories, we computed a category clustering index (CCI), which contrasts the distances of stimuli within the same category with the distances for stimuli originating from different categories. Based on the RDM computed for the 1000 test stimuli (100 stimuli per each of 10 categories), CCI contrasts distances of category exemplars within the category with distances across categories. It is defined as *CCI = (across - within) / (across + within)* and was computed for each layer of each network instance trained. CCI has a maximum of 1 (all categories cluster perfectly and are perfectly separable), and a minimum of 0 (no separability, same distances across and within categories).

In addition, we investigated the relationship between CCI and representational consistency. For each layer we computed the mean representational consistency across all 45 pairwise comparisons between 10 network instances and used Pearson correlation to demonstrate its relation to the mean class clustering indices (CCIs) across all 10 training seeds (Figure 7).

**Fig 7.**
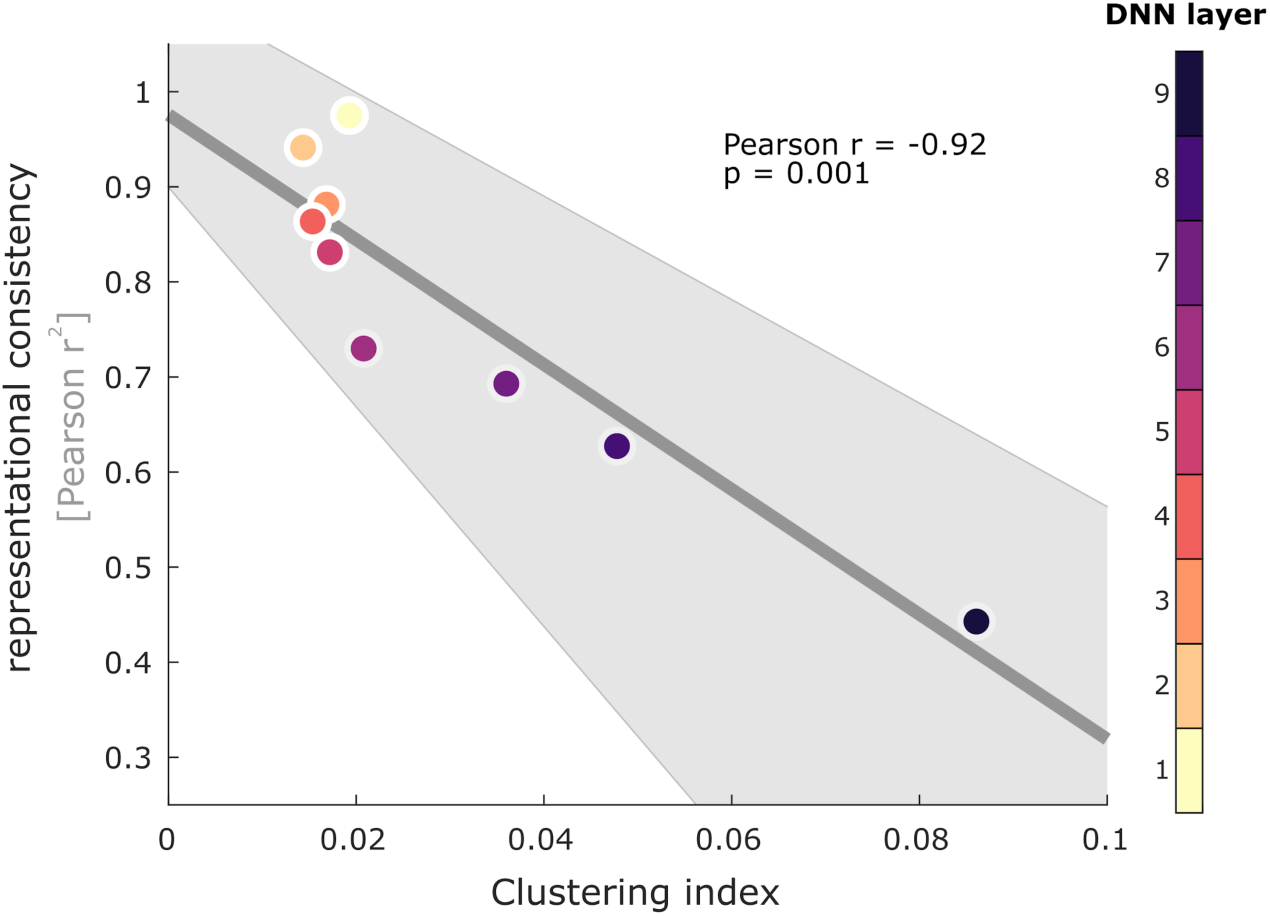
Representational consistency and category clustering are negatively correlated. Optimized for categorization performance, deep neural networks aim to separate images from different categories in the network activation space. Here we show that increasing category separability across All-CNN-C network layers (estimated here by a category clustering index) exhibits a negative relationship with the mean representational consistency across all trained network instances. Individual differences emerge while category clustering increases (95% bootstrapped CIs shown as grey area).

### Investigating causes for decreasing representational consistency

To better understand the origins of changes in representational consistency, we compare (i) exemplar-based consistency, (ii) centroid-based consistency, (iii) consistency of within-category distances, and the (iv) effects of cocktail-blank normalization.

To understand whether a misalignment in the arrangement of individual category exemplars or the arrangement of entire classes is leading to decreased consistency, we computed the 10 class centroids and used their position in activation space to arrive at centroid-based representational consistency. This was compared with consistency based on all 1,000 stimuli (exemplar-based representational consistency), and consistency computed when only distances between exemplars of the same categories were considered (within-category consistency).

To rule out effects of changed RDM size in case of centroid-based RDMs (centroid RDMs contain 45 pairwise distances whereas the exemplar-based RDMs are composed of 499,500 entries), we computed a null distribution of RDM consistency based on centroids computed from randomly sampled classes.

Finally, to test in how far the distance measure used, rather than the representational geometries themselves, could be the source of individual differences (see Supplemental Materials), we performed a cocktail blank normalization by subtracting the mean activation pattern across all images from each network unit, before computing the RDMs and representational consistency.

#### Experiments with regularization (Bernoulli dropout)

In an additional set of experiments, we explored how network regularization (here in the form of Bernoulli dropout) can affect network internal representations. Using the full CIFAR-10 set, we trained a set of 10 networks for each of 9 dropout levels (dropout probability ranging from 0 to 0.8, each of the resulting 90 DNNs was trained for 350 epochs). After training, we extracted network activations for a set of test images either by using no dropout at test time or by using multiple dropout samples for each test image. We obtained up to 10 samples extracted for each image while keeping the Dropout mask identical across network instances and the dropout rate identical to training. We created one RDM per sample and then averaged up to 10 RDMs to obtain a single RDM representing the expected representational geometry upon dropout sampling.

## Supplemental materials

**Fig S1.**
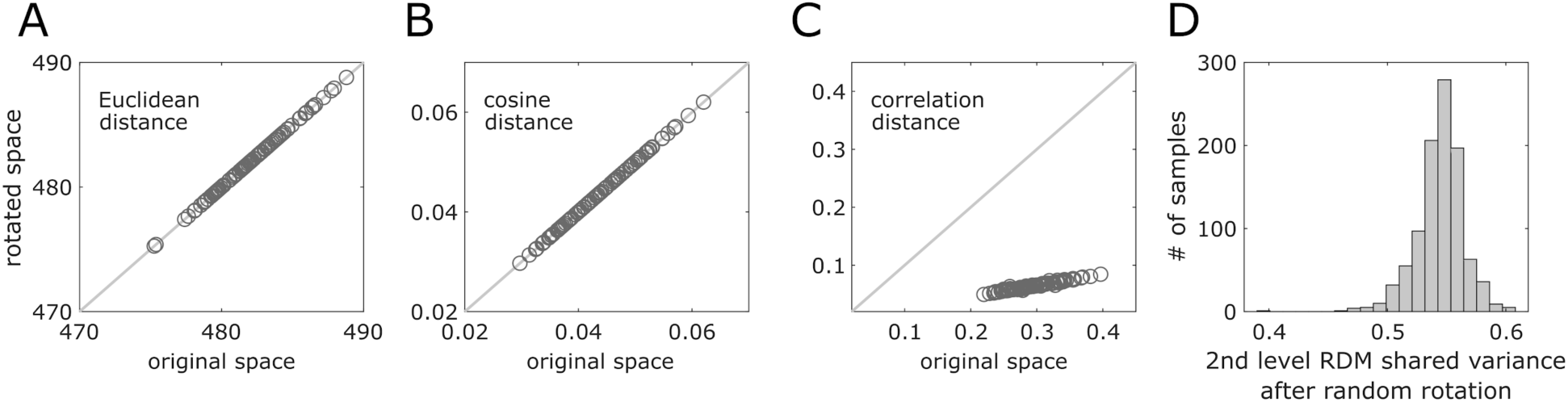
Rotation sensitivity of correlation distance. We computed the distance between two random vectors before and after both vectors were randomly rotated around the origin using the same rotation matrix. This procedure was performed for 100 vector pairs in the above simulation. Rotating both vector pairs does not have an effect when Euclidean or cosine distance is used to compute the vector pair distances (**A, B**). However, when correlation distance is used, rotations around the origin lead to decreased overall distances, and an imperfect correlation between the two distance estimates. Computing a correlation distance involves a projection of the two vectors onto a plane cutting through the origin that is orthogonal to the all-1-vector. This projection differs if the original vectors are rotated. (**C**). Accordingly, when RDMs are based on correlation distance (here based on 10 example responses), rotations around the origin lead to decreased representational consistency, despite the fact that the relative arrangement of datapoints remained identical after the rotation (**D**).

**Fig S2.**
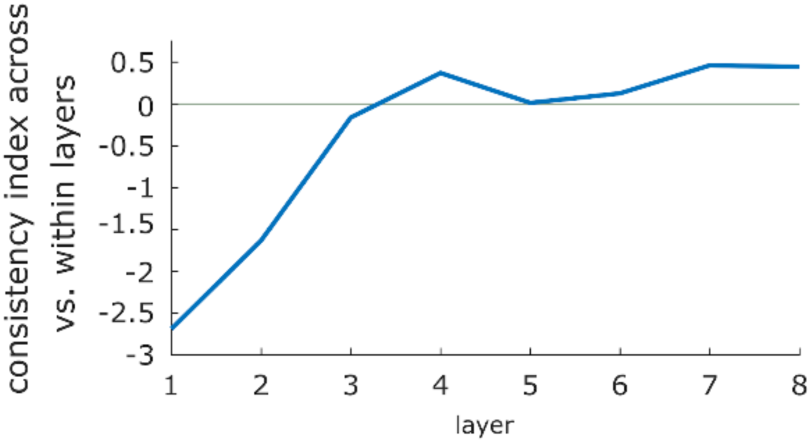
Consistency index across vs. within layers. We computed consistency across network instances (and within layers, e.g. off-diagonal elements in Fig. 3 A cell_layer_4,layer_4_) and subtracted its mean from consistency computed within instances (and across layers, e.g. diagonal elements in Fig. 3 A in cell_layer_4,layer_5_), standardized by the overall mean. This indicates that starting at layer 4, network instances are more consistent across adjacent layers than instances within layers.

